# Accumulation-depuration data collection in support of toxicokinetic modelling

**DOI:** 10.1101/2021.04.15.439942

**Authors:** Aude Ratier, Sandrine Charles

**Affiliations:** Université de Lyon, Université Lyon 1, CNRS UMR5558, Laboratoire de Biométrie et Biologie Evolutive, 69100 Villeurbanne, France

## Abstract

Regulatory bodies requires evaluation of bioaccumulation of chemicals within organisms with the objective to better assess risks linked to the toxicity of active substances. To this end, toxicokinetic (TK) data are particularly useful to relate the chemical exposure concentration to the accumulation and depuration processes happening within organisms. The bioaccumulative property of substances is quantified by bioaccumulation metrics obtained by fitting TK models to data collected from bioaccumulation tests. The internal concentrations of the studied substances are measured within organisms at regular time points during both accumulation and depuration phases, and their time course is captured by TK models thus providing bioaccumulation metrics. Still today, raw TK data remain difficult to access, as most of the time provided within papers in plots only. To increase accessibility to TK data, we present in this paper a wide collection of raw data sets extracted from the scientific literature in support of TK modelling to be performed with the MOSAIC_bioacc_ web application (https://mosaic.univ-lyon1.fr/bioacc/).

## Background & Summary

The Environmental Risk Assessment (ERA) workflow for chemicals of interest, as described in some European regulations (*e*.*g*., for plant protection products in marketing authorisation applications (EU regulation No 283/2013)), requires a bioaccumulation test, as for example on fish according to the OECD Test guideline 305^1^. Such a test consists in an accumulation phase followed by a depuration one and the time course of the internal concentration within fish is measured at regular time points during both phases. The resulting data allow to model the time course of the exposure within organisms (denoted toxicokinetic, TK), summarized in the end via bioaccumulation metrics (BCF/BSAF/BMF for water, sediment and food exposure, respectively). From a regulatory point of view, these bioaccumulation metrics are key decision criteria to determine the bioaccumulative property of active substances^2^, and to further assess potential risks that are associated to them according to the exposure sources.

Kinetic bioaccumulation metrics are always defined as ratios between uptake and elimination rates, these latter being estimated from TK models^1,3^. More precisely, TK models relate the exposure concentration to a given substance to the internal concentration within organisms, considering various processes such as accumulation, depuration, metabolism and excretion (ADME)^4^. Different types of TK models have been proposed, all being compartment models^5,6^. Organisms are usually described as unique compartments with input and output fluxes described by the TK models. Data collected from standard bioaccumulation tests are used to fit the TK models leading to uptake and elimination rate estimates. Finally, bioaccumulation metrics are calculated in support of regulatory decision making.

Some years ago, the United States Environmental Protection Agency (US EPA) has created a database for ecotoxicology data providing such bioaccumulation metrics according to species-compound combinations^7^. However, raw data are rarely provided in this database as this would require to collect them directly from the corresponding scientific papers. Moreover, when available, raw data are often provided as plots only. Nevertheless, to make updates or develop new TK modelling frameworks, it is crucial for researchers to benefit of a collection of raw TK data in order to check the robustness of their new approaches. While developing the MOSAIC_bioacc_ web application^8^ (https://mosaic.univ-lyon1.fr/bioacc/), we dealt with such a lack of raw data to fully test our innovative approach. Indeed, MOSAIC_bioacc_ provides estimates of TK model parameters and bioaccumulation metrics with their uncertainties for a large range of species-compound combinations (*e*.*g*., aquatic or terrestrial organisms exposed to metals, hydrocarbons, active substances, etc.), encompassing different exposure routes and elimination processes. For example, it was particularly difficult to collect raw TK data with biotransformation processes being involved^9^, preventing us to fully test the robustness of this tool for the widest as possible diversity of input data types.

This motivated us in creating a new public database presented in this paper. This new database gathers together more than 90 data sets of published bioaccumulation tests, concerning more than 20 genus and more than 70 substances. Some data sets concern several exposure routes (water, soil or sediment, and/or food) and several possible elimination processes (excretion, growth dilution and/or biotransformation). All the collected data were standardized in the same units and uploaded in MOSAIC_bioacc_, ensuring the use of the same methodology to get bioaccumulation metrics for all the data sets. This database should allow to overcome the current lack of raw TK data in ecotoxicology. Indeed, our purpose with this first version of the database is to motivate other researchers to share their data based on the FAIR (Findability, Accessibility, Interoperability and Reuse) principles which became today almost a duty. Our database can be considered as a proof of concept of the added-value of sharing raw TK data. It allows anyone to reuse data, for example to test new modelling frameworks. This database should also facilitate the design of new bioaccumulation experiments, as well as the comparison of results between several studies, benefiting of the unified calculation method for bioaccumulation metrics as provided by MOSAIC_bioacc_. Finally, we hope that this database could help changing of paradigm in ecotoxicology, through an increase in a more wide sharing of raw data that would certainly lead to better knowledge in the perspective of ERA.

## Methods

The conceptual workflow we followed to collect the raw TK data is summarized on Figure 1 and detailed below.

**Figure 1.**
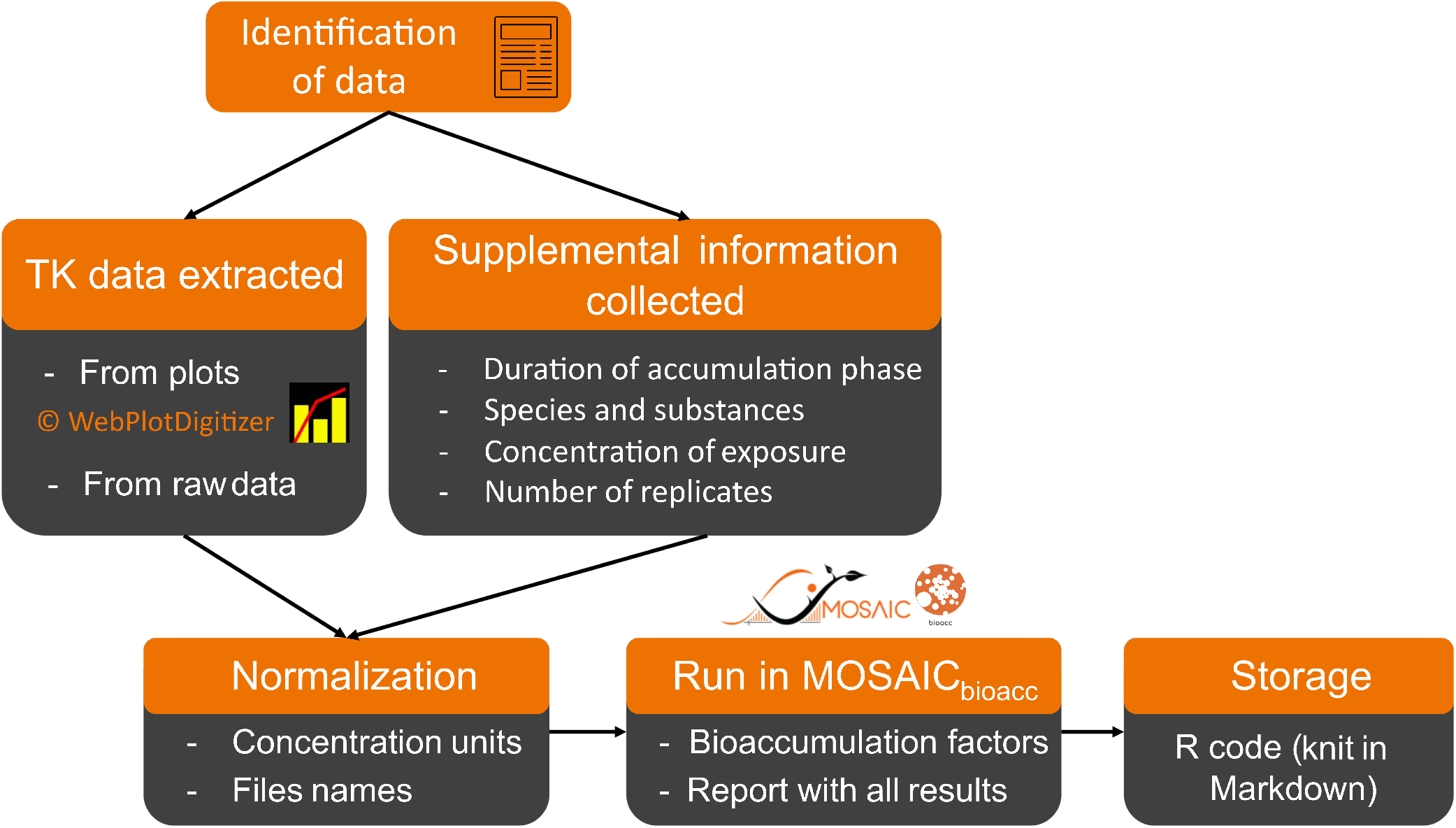
Conceptual framework of the collection of TK data and their storage in the database of MOSAIC_bioacc_.

### Data and literature sources

One of our aims was to test the robustness of the generic feature of the MOSAIC_bioacc_ methodological framework from a wide collection of TK data sets, encompassing a variety of genus, substances, exposure routes and elimination processes. For this purpose, the US EPA ecotoxicology database^7^ could not be used as raw observations are usually absent together with a lack of information (*e*.*g*., biotransformation processes are usually not informed). Consequently, an in-depth search in literature with Scopus was performed with specific keywords according the expected type of TK data sets. These keywords were for example: “TK model aquatic”, “TK model terrestrial”, “TK model biotransformation”, “TK biotransformation rate”, “TK model food exposure”, “TK model sediment exposure”, “TK model water exposure”, etc. For each article retained, we manually checked if TK data were available in a table or in a plot, in the manuscript or in the supporting information. A total of 28 studies was finally selected for a full analysis with a TK model.

### Toxicokinetic data

#### Literature source data extraction

Only few publications provided raw data as table of internal concentrations of a chemical (and its potential metabolite(s)) within organisms versus time. For most of the studies, data were only available in plots. Thus, plots were converted in JPEG files from screenshots (native screenshot tool in Windows 10). Then images were imported in WebPlotDigitizer^10^, which assists the users to extract the underlying numerical data of a plot. The calibration of axes and the selection of points for extraction is performed by the user. Raw data are exported from WebPlotDigitizer to CSV files. Then, CSV files data were formatted for MOSAIC_bioacc_ requirements, manually adding information on replicates and exposure concentration according to the Methods section of each article. For each collected TK data set, the corresponding file was named by the genus, the substance, the duration of the accumulation phase, the first author and the year of publication, possibly adding comments (*e*.*g*., size of the tested organism) when available.

#### Data standardization

Data were standardized to always express exposure concentrations in *µg*.*mL*^*–*1^ (for water exposures) or in *µg*.*g*^*–*1^ (for sediment or food exposures). The internal chemical concentrations in organisms were translated in *µg*.*g*^*–*1^, as required for a proper use with MOSAIC_bioacc_.

#### MOSAIC_bioacc_ running

Once standardized, each collected data set was uploaded in MOSAIC_bioacc_ (https://mosaic.univ-lyon1.fr/bioacc). The generic modelling framework of MOSAIC_bioacc_ automatically provided both kinetic and steady-state bioac-cumulation metrics, unifying their calculation among data sets allowing relevant comparisons or a correct classification of substances according to their bioaccumulative property for example. Kinetic bioaccumulation metrics (given as medians and 95% credible intervals) were saved in a local repository, and the report with all fitting outputs downloaded in the same repository, for each data set.

### Storage

An R project^11^ was created in a local repository of the MOSAIC_bioacc_ Shiny application, containing all collected data sets, the bibtex file with all references, all reports and all kinetic bioaccumulation metric estimates. Then, an RMarkdown file^12,13^ allows to create the final table based on information collected from the name of the data set and from the data set itself (*e*.*g*., column headers, number of data, number of replicates), as well as from the bibtex file. The R package ‘DT’^14^ was used to combine all collected information in a user-friendly table, and the RMarkdown file was knited in HTML format for display to the user. In the future, each new data set added into the repository will make the RMarkdown automatically knited^13^ for update.

## Data Records

The database can be accessed directly on-line at http://lbbe-shiny.univ-lyon1.fr/mosaic-bioacc/data/database/index_readme.html or via MOSAIC_bioacc_ clicking the “More scientific TK data” button. An example of the output of the database is shown in Figure 2. The collected raw TK data of the database consist in time courses of several types of chemicals bioaccumulated in various species via different exposure routes. Each data set is summarized by:

**Figure 2.**
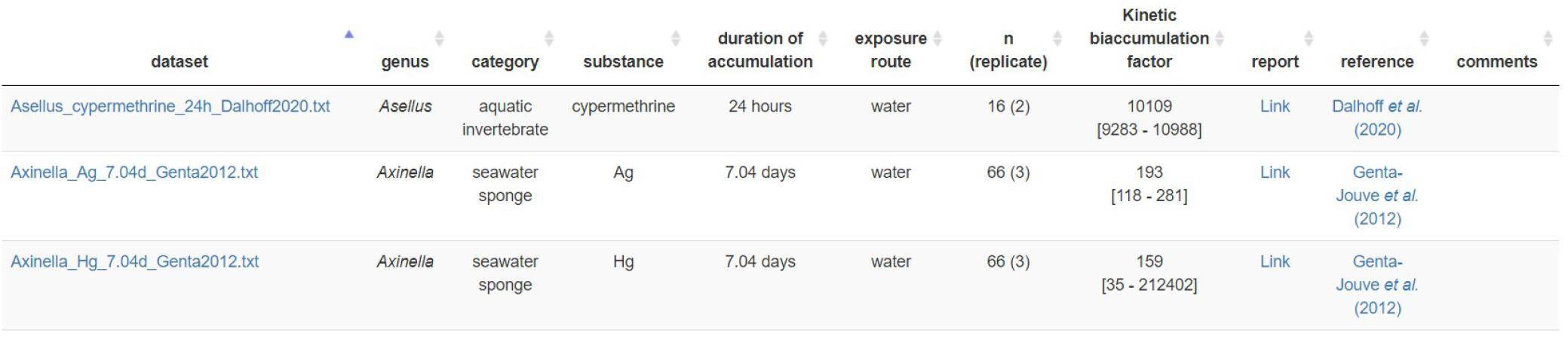
Screenshot of a sample from the database available from MOSAIC_bioacc_.

- the file name (raw data directly downloadable by clicking on the file name, in TXT or CSV format),
- the genus of the tested organism,
- the category of the organism (*e*.*g*., aquatic, terrestrial, etc.),
- the tested substance,
- the duration of the accumulation phase,
- the tested exposure routes and for which data are available (*e*.*g*., water, sediment, food, pore water),
- the total number of data in the data set (and in brackets the number of replicate(s)),
- the kinetic bioaccumulation metric median value with its 95% credible interval,
- the report which contains all the results from MOSAIC_bioacc_ (in PDF file),
- the link to the reference or the source of data,
- some additional comments (*e*.*g*., lipid fraction, growth, biotransformation).

A summary of all data sets is presented in Table 1. Genus were separated in nine categories: aquatic invertebrates (*n* = 51), fish (*n* = 21), insects (*n* = 5), aquatic worms (*n* = 5), terrestrial worms (*n* = 3), seawater sponges (*n* = 2), seawater plants (*n* = 1), aquatic algae (*n* = 1) and terrestrial invertebrates (*n* = 1). The most represented genus in the database are *Oncorhynchus* (fish) and *Gammarus* (aquatic invertebrate), followed by *Daphnia* (aquatic invertebrate), classically used in ecotoxicological experiments. Recommended genus by OECD guidelines for bioaccumulation tests are *Eisenia* and *Enchytraeus* for terrestrial organisms (OECD 317)^15^, and *Tubifex* or *Lumbriculus* for aquatic invertebrates exposed to sediment (OECD 315)^16^; data sets for these species are available in the database (*n* = 8).

**Table 1.** Summary of collected data. The classification of bioaccumulative properties are the same as in ECHA (2017)^2^.

| Genus (n = 25) | Substances (n = 64) | Exposure route | Elimination process | Kinetic bioaccumulation metric (median) | Number of studies |
| --- | --- | --- | --- | --- | --- |
| aquatic invertebrate (n = 51)<br>fish (n = 21)<br>insect (n = 5)<br>aquatic worm (n = 5)<br>terrestrial worm (n = 3)<br>seawater sponge (n = 2)<br>seawater plant (n = 1)<br>aquatic algae (n = 1)<br>terrestrial invertebrate (n = 1) | metals (n = 9)<br>PCB (n = 5)<br>pesticides (n = 23)<br>flame retardants (n = 3)<br>pharmaceutical products (n = 2)<br>hydrocarbons (n = 14)<br>octyphenols (n = 1)<br>polyesters (n = 2)<br>nanoparticles (n = 5) | water (n = 57)<br>food (n = 21)<br>sediment (n = 12) | biotransformation (n = 18)<br>growth dilution (n = 0) | < 1000 (n = 75)<br>1000 < BAF < 2000 (n = 4)<br>2000 < BAF < 5000 (n = 3)<br>> 5000 (n = 8) | 28 |

Substances were divided in nine classes: pesticides (*n* = 23), hydrocarbons (*n* = 14), metals (*n* = 9), nanoparticules (*n* = 5), polychlorobiphenyls (PCB, *n* = 5), flame retardants (brominated or chlorinated, *n* = 3), pharmaceutical products (*n* = 2), polyesters (*n* = 2) and octyphenol (*n* = 1). Among all data sets, the majority of bioaccumulation tests were performed via spiked water. Besides, 18 data sets account for biotransformation processes, considering from 1 to 8 metabolites.

According to ECHA (2017)^2^, bioaccumulation metrics below 1,000 mean that the substance is not bioaccumulative, whereas one ranging between 2,000 and 5,000 corresponds to a bioaccumulative substance. If the bioaccumulation metric is higher than 5,000, the substance is classified as very bioaccumulative. These ranges are reported in Table 1, where bioaccumulation metrics are higher than 5,000 for eight data sets, indicating the very bioaccumulative property of the substances for the studied genus. On an ecotoxicological point of view, the most higher bioaccumulation metrics were obtained for *Daphnia* exposed to alpha-cypermethrin, triphenyltin hydroxide (TPT), diazinon and imidacloprid; *Hydropsyche, Chaetopteryx, Tubifex* and *Asellus* exposed to cypermethrin, highlighting the potential toxicity of cypermethrin for aquatic and terrestrial organisms.

## Technical Validation

The validity of the collected data was checked by considering several aspects in the corresponding papers: (i) limitations in the method and/or source material, (ii) unreported replicates (experimental variability missing), (iii) inaccuracy in the collection techniques used to extract data from source content, and (iv) the quality of the results we obtained (based goodness-of-fit and precision criteria) when reusing the data in our TK modelling framework.

For some studies, raw data were not always consistent with the associated plot (*e*.*g*., differences in time points or measured concentrations in organisms) as provided within the original paper. When a possible inaccuracy collection of data was noticed, the data set was removed from the database. Besides, most of the studies did not provide all data, for example the measurement concentration in each replicate were missing. Indeed, most of the time, plots only provide average measured data with error bars representing standard deviation, standard error or other uncertainty measures, without giving access to the raw observed concentrations. In such cases, only the mean values were collected.

The uncertainty due to data extraction from plots with the WebPlotDigitizer tool^10^ is legitimate to consider in order to see if a bias was introduced in collected data sets. To quantify this bias, we performed ten times the collection of TK data for a same data set (.jpeg file) and calculated the extraction errors possibly due to the shape of points, the axis calibration, the image resolution or some other factors. For example, a variation of 0.12 *±* 0.13% was observed for the data set of Chen *et al*., (2017)^17^ (Figure 2-a in their paper), which is relatively low (*<* 0.5% as usually considered for coefficients of variation). This is representative of what we had for most of the plots.

In order to support the quality of each data set provided in our database, MOSAIC_bioacc_ was used to fit TK models for all data sets under the same conceptual framework. Hence, the bioaccumulation metrics and TK model fitting plots are systematically provided (Figure 3), as well as a collection of goodness-of-fit (GOF) criteria allowing to check the relevance of the results. All this information is gathered together in a PDF report downloadable from the MOSAIC_bioacc_ database.

**Figure 3.**
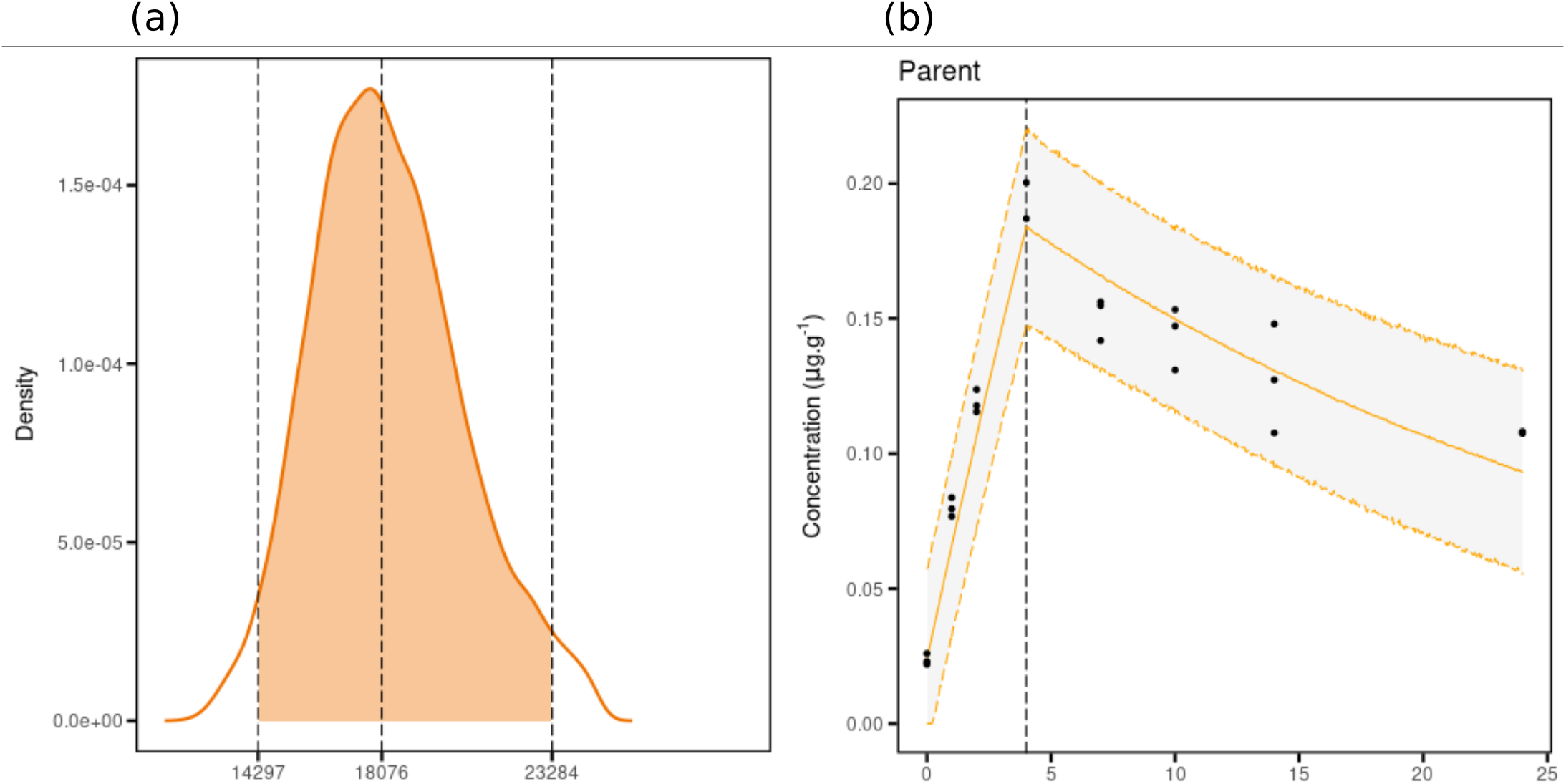
Example of results given by MOSAIC_bioacc_: (a) the bioaccumulation metric and (b) the fitted TK model predictions with observed data. This example comes from the data set ‘Male_Gammarus_Single.txt’ available from MOSAIC_bioacc_.

More than 95% of data sets of the MOSAIC_bioacc_ database are already published. The reuse with MOSAIC_bioacc_ allows to compare fitting results to those of the original study and to confirm the relevance of including the data in the database. In addition, most of the publications (25 among 28) do not provide uncertainty around predicted concentrations (at least not visualized in the original plots), while we provide, for each data set, the 95% credible intervals around median predicted concentrations, as well as uncertainties for all bioaccumulation metrics. It is an undeniable added-value we bring to the ecotoxicological community thanks to the revisit of all these data sets.

## Usage Notes

Raw TK data are particularly useful to better understand the relationship between exposure chemical concentrations in the environment and their potential bioaccumulation within organisms. They are also used by researchers or regulators to evaluate the bioaccumulative property of chemicals then to establish the link with the life-history traits of effects on organisms. Our database also gives an easy access to formatted raw TK data that can be used to improve current TK models or to develop new conceptual modelling frameworks. This database is not an overview of all available TK data in the literature, but it encompasses several types of genus, substances, routes of exposure and elimination processes, giving a proof of concept for raw TK data storage and sharing. Our database aimed at developing and validating our generic TK modelling framework, as illustrated in this paper where the evaluation of the robustness of MOSAIC_bioacc_ through the whole collection of data sets proved to be successful. In the past, evaluation and improvement of TK modelling frameworks have been limited due to the lack of raw TK data for a large variety of species and chemicals, especially for data considering biotransformation^9^, often more scarce than classic TK data. From our database, such evaluation and improvement should be facilitated and researchers are invited to contribute to this database by sharing their own data sets.

If bioaccumulation testing is required, it may be difficult to plan the experimental design since its depends on the substance, the species and the exposure route under consideration. Usually, a search in literature provides a possible range of exposure concentrations for the substance, but the amount likely to be bioaccumulated in organisms of the studied species remains unknown, as well as how long it will require to reach the steady-state at the end of the accumulation phase. Hence, the MOSAIC_bioacc_ database provides a quick search option to identify similar genus, or to choose a substance with a similar mode of action to consider in support of new experiments to plan.

We sincerely hope this database will support further research to increase our knowledge on bioaccumulation of substances for which there are today clear societal challenges of environmental protection. Indeed, in ERA, bioaccumulation metrics are key decision criteria to classify a substance as non-bioaccumulative, bioaccumulative or very bioaccumulative. Most often, these metrics are calculated from two or three different modelling approaches, which makes their comparison difficult, in addition to be given only as a point mean value (without uncertainty). Our database should then improve this classification thanks to the kinetic bioaccumulation metrics we provide from the same methodology for all data sets, that clearly quantify the uncertainty summarized by the 95% credible intervals around median values. More globally, our database should strongly encourage researchers to share their data with the scientific community by directly providing the raw data in their publications.

## Code Availability

Data sets are available at MOSAIC_bioacc_ (https://mosaic.univ-lyon1.fr/bioacc) or directly at http://lbbe-shiny.univ-lyon1.fr/mosaic-bioacc/data/database/index_readme.html. To reproduce all the outputs provided in the reports for all data sets of the database, data can be downloaded from the database then uploaded in MOSAIC_bioacc_. In addition, for each data set, the corresponding R code can be downloaded if using the R software directly is the preferred option of the user.

## Acknowledgements

The authors are thankful to ANSES for providing the financial support for the development of the MOSAIC_bioacc_ web tool. This work was performed using the computing facilities of the CC LBBE/PRABI. This work benefited from the French GDR “Aquatic Ecotoxicology” framework which aims at fostering stimulating scientific discussions and collaborations for more integrative approaches. This work is part of the ANR project APPROve (ANR-18-CE34-0013) for an integrated approach to propose proteomics for biomonitoring: accumulation, fate and multi-markers (https://anr.fr/Projet-ANR-18-CE34-0013). This work was also made with the financial support of the Graduate School H2O’Lyon (ANR-17-EURE-0018) and “Université de Lyon” (UdL), as part of the program “Investissements d’Avenir” run by “Agence Nationale de la Recherche” (ANR). The authors are also thankful to Nina Ceredergreen, Frédéric Gimbert and Roman Ashauer to their help in providing raw TK data.

## Author contributions statement

A.R. and S.C. participated to the collection of data sets, analysed the results in MOSAIC_bioacc_ and elaborated the database. All authors reviewed the manuscript.

## Competing interests

The authors declare no competing interests.

## Notes

http://lbbe-shiny.univ-lyon1.fr/mosaic-bioacc/data/database/index_readme.html

### Competing Interest Statement

The authors have declared no competing interest.

